# Sequencing smart: *De novo* sequencing and assembly approaches for non-model mammals

**DOI:** 10.1101/723890

**Authors:** Graham J Etherington, Darren Heavens, David Baker, Ashleigh Lister, Rose McNelly, Gonzalo Garcia, Bernardo Clavijo, Iain Macaulay, Wilfried Haerty, Federica Di Palma

## Abstract

**Background:** Whilst much sequencing effort has focused on key mammalian model organisms such as mouse and human, little is known about the correlation between genome sequencing techniques for non-model mammals and genome assembly quality. This is especially relevant to non-model mammals, where the samples to be sequenced are often degraded and low quality. A key aspect when planning a genome project is the choice of sequencing data to generate. This decision is driven by several factors, including the biological questions being asked, the quality of DNA available, and the availability of funds. Cutting-edge sequencing technologies now make it possible to achieve highly contiguous, chromosome-level genome assemblies, but relies on good quality high-molecular-weight DNA. The funds to generate and combining these data are often only available within large consortiums and sequencing initiatives, and are often not affordable for many independent research groups. For many researchers, value-for-money is a key factor when considering the generation of genomic sequencing data. Here we use a range of different genomic technologies generated from a roadkill European Polecat (*Mustela putorius*) to assess various assembly techniques on this low-quality sample. We evaluated different approaches for *de novo* assemblies and discuss their value in relation to biological analyses.

**Results:** Generally, assemblies containing more data types achieved better scores in our ranking system. However, when accounting for misassemblies, this was not always the case for Bionano and low-coverage 10x Genomics (for scaffolding only). We also find that the extra cost associated with combining multiple data types is not necessarily associated with better genome assemblies.

**Conclusions:** The high degree of variability between each *de novo* assembly method (assessed from the seven key metrics) highlights the importance of carefully devising the sequencing strategy to be able to carry out the desired analysis. Adding more data to genome assemblies not always results in better assemblies so it is important to understand the nuances of genomic data integration explained here, in order to obtain cost-effective value-for-money when sequencing genomes.

## Introduction

Starting in 1990, the Human Genome Project used low-throughput, high-cost Sanger sequencing platforms to create the first draft human genome at a cost of USD $300 million. Fast-forward 19 years and the cost of sequencing a human genome has dropped to around USD $1000. Short-read technologies producing high-throughput, low per-base cost next-generation sequencing (NGS) means that genomics is no longer restricted to large sequencing consortiums and has opened up the field to even the smallest of research groups. The recently formed Vertebrate Genomes Project (VGP) [1] aims to produce near-gapless, chromosome-scale phased genome assemblies for around 66,000 extant vertebrate species. The assembly pipeline consists of 60x coverage Pacific Biosciences (PacBio) long read sequencing, followed by 10x Genomics linked reads, Bionano optical mapping and Arima Genomics’ Hi-C profiles. These long-read technologies provide highly contiguous genome assemblies. Similar consortiums and sequencing initiatives have been formed to sequence a range of target organisms such as Bat1K, Bird10K, Oz Mammals Genomics, and Earth BioGenome Project (including Darwin UK Tree of Life, Colombia EBP, etc.) [2]. Although such efforts make it possible to achieve highly-contiguous, chromosome level genome assemblies, the cost of generating this amount of data and assemble them is considerable and often only within reach of a few of these consortiums. It is important for smaller independent research groups or initiatives to consider value-for-money against biological questions as a key factor when planning the generation of genomic sequencing data.

### Non-model organism

Non-model organisms have the potential to provide new knowledge related to phenotypic and genotypic variation. Through comparative genomics, it is possible to identify how different organisms are related to each other, how they adapt to novel environments, or the genetic basis underlying novel phenotypes. These new findings can be applied to further research, such as in the biomedical and food industries through breeding programs with the development of marker assisted selection and in conservation biology [3-12] *De novo* assembly of endangered species, followed by low-coverage population-level sequencing provides unprecedented information about the amount of genetic diversity within populations, past and ongoing gene flow between different populations, and the level of inbreeding in small populations.

However, there are a number of difficulties when working with non-model mammals. Firstly, the genome size is not always known, hampering the assessment of the completeness of the ‘assembled’ genome and of the sequencing depth. Additionally, the availability and quality of the samples used for sequencing non-model organisms is often substandard. Tissue and blood samples are often obtained from wild populations and may need to be acquired from remote locations, delaying the time between collection and DNA extraction. Another common issue relates to samples which may have been stored in collections such as museums, zoos and tissue collections and subjected to a number of different preservation methods such as freezing, storage in ethanol, FFPE, etc. Many current sequencing technologies (e.g. PacBio, Bionano, and 10x Genomics) rely on high-molecular-weight DNA with molecules longer than 100Kb being optimum. Degraded DNA, as is commonly observed in samples from wild populations and is usually sub-optimal for use in many advanced sequencing methods. It is therefore difficult, or sometimes impossible, to leverage the full application of these technologies.

Many non-model organisms are species from wild populations that are highly heterozygous leading to numerous challenges during the assembly step. Allelic differences in a diploid genome generates branches and bubbles in the assembly graph [13]. Even though most graph-based assemblers have functions to search for and remove these structures, high density variation can still make assembly of heterozygous organisms challenging. Conversely, high levels of homozygosity, characteristic of endangered (and typically inbred) species, hamper the efforts of creating phased genome assemblies, since the ability to phase haplotypes is dependent on linked sequences spanning polymorphisms. Additionally, non-model organisms vary in their ploidy, chromosome number, repeat content, sequence composition and GC content, adding further confounding factors to genome assembly.

### The example of the European Polecat

The European Polecat (*Mustela putorius*) is a medium-sized carnivore found across Europe and the Middle East. It is purported to be the ancestral species of the domestic ferret (*M. p. furo*) [14]. Across most of mainland Europe the polecat is in widespread decline [15]. In the United Kingdom, the European Polecat has a chequered history. Persecuted to the verge of extinction in the early-1900’s, when it was confined to unmanaged forests in central Wales, it has since seen a population increase and is now found throughout Wales and across much of central, south-western and eastern England [16].

Here, a road-kill sample of European Polecat from the Vincent Wildlife Trust collection (VWT 693) was used to assess short-read and long-range *de novo* sequencing strategies for non-model mammals. Comparisons between combinations of PCR-free Illumina libraries, Nextera long-mate-pair (LMP) libraries, 10x Genomics Chromium libraries and Bionano optical maps are made to assess optimum sequencing and assembly strategies.

#### Sequencing technologies

##### Short-read sequencing

The market-leader in short-read high-throughput NGS is Illumina [17]. Recent machines produce read-lengths of 100 bps and above and a single Illumina Novaseq run is currently capable of generating 600 Gbps of read data. An advantage to Illumina sequencing is the generation of paired-end (PE) reads, in which the sequence from both ends of each DNA molecule is synthesised. As the input molecules are of an approximate known length, the acquisition of PE data provides a greater amount of information. Additionally, using a PCR-free library preparation removes bias in genomic coverage, previously incorporated by a PCR amplification step in older library preparation procedures. PCR-free Illumina sequencing requires a minimum of 2-5 µg of genomic DNA (gDNA) at a minimum concentration of 35 ng/µl in 60 µl.

##### Long Mate Pair sequencing

Long DNA fragments up to around 40 kb can be sequenced to provide PE reads that bridge long repeats, thus producing longer contiguous genome assemblies as well as characterising structural variants. Under the Nextera LMP protocol [18], a transposase enzyme attaches 19-bp biotinylated adaptors to both ends of each long DNA fragment. The DNA is then circularized, where the biotinylated ends become joined. The circularized DNA is then fragmented and biotin enrichment is used to process the fragments containing the adaptors that mark the junction. During sequencing, reads are produced from both ends of a fragment, resulting in inward-facing reads that read toward and through the adaptors. For Illumina Nextera LMP sequencing the Nextclip tool can then be used to trim adaptors and de-duplicate reads [19]. Nextera LMP sequencing requires a minimum of 9 µg of gDNA at a minimum concentration of 30 ng/µl in 300 µl.

##### 10x Genomics

The Chromium system from 10x Genomics uses oil emulsion and multiple displacement amplification (MDA) to ligate short molecular barcodes to reads from each fragment of DNA [20], followed by PE Illumina sequencing. Each fragment receives its own unique barcode and hence reads with the same barcodes represents clusters of reads from the same region in the genome. These ‘linked-reads’ provide the long-range information missing from standard Illumina sequencing and is then used to assemble phased assemblies *de novo*. 10x Chromium libraries require high molecular weight gDNA at a concentration of 20µg/µl in 10µl. gDNA should greater than 50kb in length in order to take full advantage of the technology.

##### Optical mapping (Bionano)

Bionano technology produces optical maps of nicking/restriction enzyme sites across kilobase-long stretches of DNA molecules, providing a high-throughput tool for ordering and orienting contigs of physical maps and validation of genome assemblies [21]. Bionano optical maps can be compared to *in silico* restriction maps produced from an NGS genome assembly for validation purposes, to improve contiguity by assigning the shorter NGS scaffolds to the longer optical maps, and identifying structural variants. 600 ng of raw gDNA at a concentration of 35-200 ng/µl is typically enough DNA to generate about 120 µl of labelled molecules – enough to provide adequate coverage for analysis of a human-sized genome (3 Gb).

Genome contiguity has an effect on what analyses can be achieved (Table 1), so it is important to appreciate the power and limitations of each sequencing strategy and technology.

**Table 1.** Information regarding the possible resolution for various *de novo* genome sequencing technologies. When planning a genome assembly project, it is important to understand the strengths and limits of the various sequencing strategies available.

| Assembly resolution | Paired-end | Paired-end +<br>Long Mate Pair | Bionano | 10x<br>Genomics |
| --- | --- | --- | --- | --- |
| Gene content | Yes | Yes | No | Yes |
| Gene order | Yes | Yes | No | Yes |
| Repeat spanning | No | Yes | Yes | Yes |
| Structural variants | No | Yes | Yes | Yes |
| Haplotype resolution | No | No | No | Yes |

## Materials and Methods

### Sequencing

Using the same sample of a roadkill European Polecat sample stored in 100% ethanol, two lanes of PCR-free Illumina HiSeq2500 250bp PE reads (77x coverage), two Illumina LMP libraries of size 5 kb (27x coverage) and 7 kb (9x coverage) and four lanes of 150bp PE 10x Genomics Chromium using an Illumina HiSeq2500 (85x coverage) are generated.

The mean molecule length of the European Polecat sample was around 50kb, which was not of good enough quality to generate Bionano data (recommended >100kb). Because the domestic ferret and its polecat ancestor diverged only around 2000 years ago, and fully interbreed we do not expect significant divergence and structural differences between the two species. Therefore, the original sample used for the domestic ferret genome assembly [22] was obtained and one chip of Bionano Genomics optical genome maps was generated. This was used to create Bionano hybrid-scaffold assemblies for the European Polecat genomes assembled with the previously mentioned short-read data. We generated 664 Gb of Bionano molecules, with an N50 size of 185 kb and a contig coverage of 261x. Of this, 40% of the molecules aligned back to the Bionano *de novo* assembly, leaving an effective coverage of 110x. A more detailed description of the library preparation methods can be found in the Supplementary Methods.

### Assemblies

10 different genome assemblies were generated as summarised in Figure 1, (with additional information in Supplementary Table S1), and detailed as follows:

**Figure 1.**
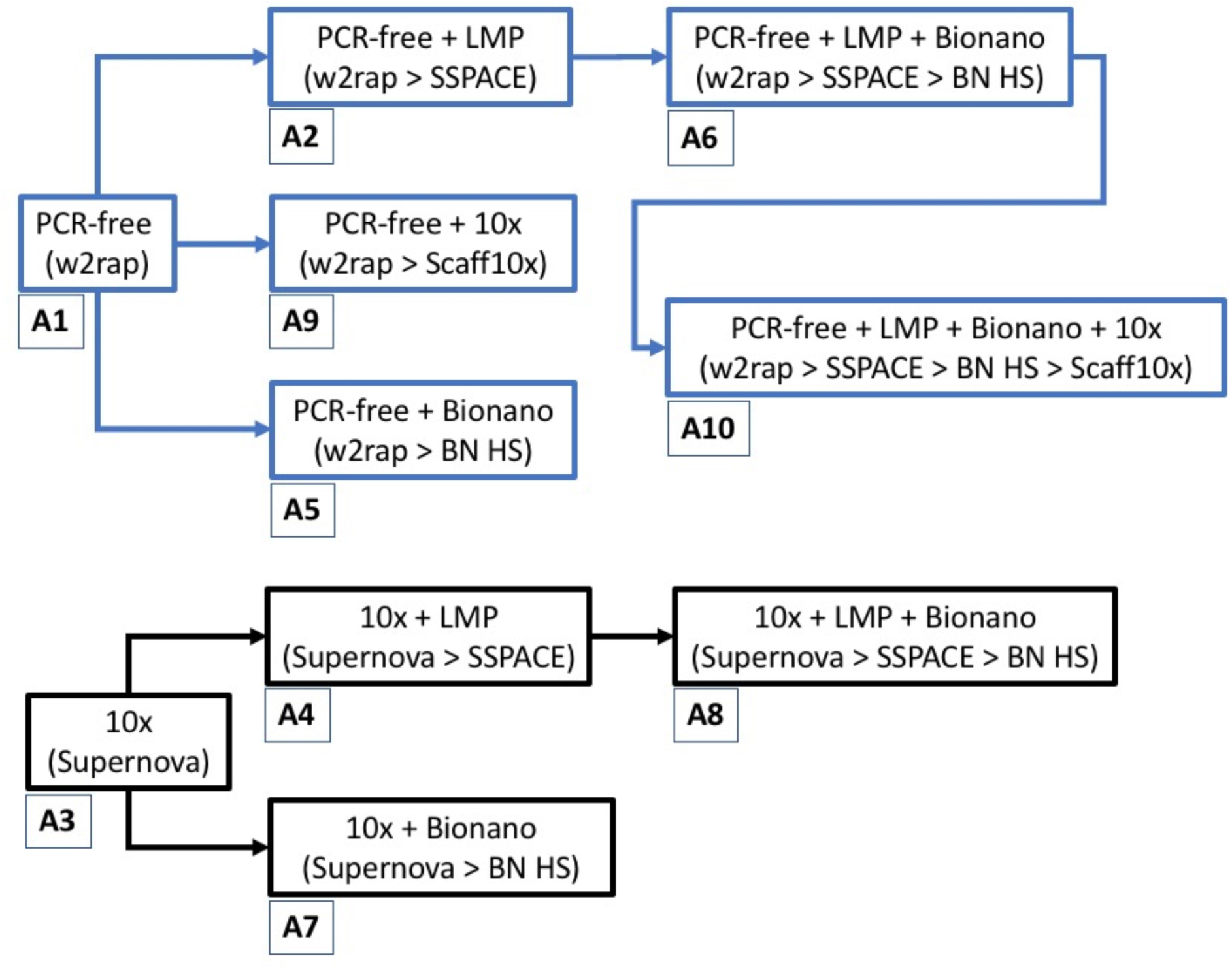
Ten different assembly strategies using a variety of different data types: PCR-free Illumina short-read (‘PCR-free’), long mate-pair (‘LMP’), 10x Genomics Chromium library (‘10x’), and Bionano Genomics optical maps (‘Bionano’). The blue-boxed assemblies all originate from the same PCR-free w2rap assembly (A1) and the black-boxed assemblies all originate from the same 10x Genomics Supernova assembly (A3). Information in brackets refers to assembly software pipeline and assembly numbers are annotated below each assembly.

#### Assembly A1 (w2rap)

The PCR-free Illumina reads from polecat were assembled using the w2rap-contigger [23]. w2rap is predominantly a contig assembler-reads are used to construct an assembly graph which is then traversed to create a contig assembly. A final step involves using the PE information to scaffold contigs not joined during the initial assembly process. Using w2rap, four different assemblies were created using a range of k-mers (k=180, 200, 224, and 240) and simple assembly stats were run to examine contiguity across the assemblies (for all contigs and filtering for contigs > 1kb). From these statistics, the assembly constructed with k=224 was selected as the final assembly.

#### Assembly A2 (w2rap + lmp)

Using SSPACE [24] the 5 kb and 7 kb Nextera LMPs were used to scaffold the w2rap assembly from assembly A1. For all SSPACE LMP assemblies the reads were used only for scaffolding and not for contig extension.

#### Assembly A3 (10x)

The 10x Genomics Chromium library were assembled using the 10x Genomics Supernova software [20], with default parameters. Similar to w2rap, Supernova creates an initial contig assembly but then scaffolds using the molecule-specific barcode information in the reads to join contigs known to be from the same molecule [20]. The output style of the resulting assembly was ‘pseudohap’, which creates one haplotype per scaffold.

#### Assembly A4 (10x + lmp)

The 5 kb and 7 kb Nextera LMPs were used to scaffold the 10x assembly generated in assembly A3. As in assembly A2, the LMP reads were used only for scaffolding and not for contig extension.

The Bionano data was assembled *de novo* and then was used to position and orient scaffolds from previous assemblies creating a Bionano hybrid-scaffold as follows:

#### Assembly A5 (w2rap + bionano)

Bionano hybrid-scaffolding with w2rap assembly (Assembly A1).

#### Assembly A6 (w2rap + lmp + bionano)

Bionano hybrid-scaffolding with the w2rap + LMP assembly (Assembly A2)

#### Assembly A7 (10x + bionano)

Bionano hybrid-scaffolding with the 10x assembly (Assembly A3).

#### Assembly A8 (10x + lmp + bionano)

Bionano hybrid-scaffolding with the 10x + LMP assembly (Assembly A4).

Finally, the 30x coverage of 10x Genomics data (from the same data generated for assembly A3, henceforth referred to in the text as ‘10x-scaffolding’) was used to scaffold two assemblies using the scaff10x program from Phusion2 [25], as follows:

#### Assembly A9 (w2rap + 10x)

The w2rap-only assembly (Assembly A1) with 10x-scaffolding.

#### Assembly A10 (w2rap + lmp + bionano + 10x)

The w2rap + LMP + Bionano assembly (Assembly A6), with 10x-scaffolding.

### Analyses

#### Genome contiguity

For each genome assembly, a number of assembly statistics, such as contig N50, scaffold N50, the number of scaffolds over given lengths and scaffolded genome size were calculated. To calculate contig N50, any scaffolded-contigs that were joined by 25 or more Ns were broken. The percentage of the genome contained in scaffolds greater than 25 kb (the average length of a vertebrate gene [26]), and the number of scaffolds greater than 47 Mb (the length of the smallest human chromosome), were also calculated

#### K-mer analysis

The K-mer Analysis Toolkit (KAT) version 2.3.4 [27] was used to examine k-mers across reads and assemblies. KAT enables users to assess levels of errors, bias and contamination at various stages of the assembly process. Using the KAT ‘comp’ program with a k-mer size of 31, k-mers in the PCR-free Illumina reads were compared with those in the resulting assemblies (omitting the Bionano assemblies as this technology adds negligible sequence content) and for each assembly, the k-mer-spectra was plotted.

#### Gene content

BUSCO (v2.0.1) was used to search for single-copy orthologs in each assembly [28]. BUSCO reports the number of single-copy orthologs discovered in the input assembly, and categorises them as ‘complete’, ‘single-copy’, ‘multi-copy’ or ‘fragmented’. For speed, 27 sequences that had tblastn runtimes of over 3 days were removed from the mammalia_odb9 database, leaving us with a final database of 4077 single-copy orthologs. This custom version of mammalia_odb9 was used for the ‘lineage’ parameter in BUSCO and ‘human’ for the Augustus species parameter.

#### Repeat content

To examine repeat content and compare how repeats were resolved in each genome assembly RepeatMasker [29] was used to identify repeat families in each assembly, using all *Carnivora*-specific repeats. As well as identifying repeat sequences, the mean deletion, insertion and divergence for each family was also calculated, as well as the mean values overall. Mean divergence is calculated as ‘mismatches/(matches + mismatches)’ between queries and matches for all repeats.

#### Assembly errors and misassemblies

REAPR [30] was used to evaluate the accuracy of each genome assembly by separately mapping PCR-free PE and LMP reads back to each assembly. The fragment coverage distribution (FCD) error for each assembly was calculated. FCD is the fragment depth from only the reads that are mapped to a given base of a fragment. The FCD error is the difference between the theoretical and observed FCD and is used to identify assembly errors in the regions containing a run of high FCD errors. Mapping information such as the FCD and insert size distribution is analysed to locate misassemblies as well as more local per-base accuracies. The ‘smalt map’ option in REAPR was used, which uses SMALT [31] to align the PCR-free PE and LMP reads back to each assembly utilising the option to map PE reads independently. This ensures that read pairs are not artificially forced to map as proper pairs within a given insert size. REAPR was then used to identify perfectly and uniquely mapped reads in the PE PCR-free alignment, to accurately call error-free bases in the assembly and further used the LMP reads to identify features consistent with misassemblies. Error-free bases have at least 5X perfect and unique coverage of paired end reads. REAPR summary scores were calculated for each assembly by multiplying the number of error-free bases with the square of the REAPR broken scaffold N50 length, and then dividing by the original scaffold N50, i.e. ‘No. error-free bases * (broken N502/assembly N50)’. This test was first used to evaluate genome assemblies in the Assemblathon series [26] and rewards local accuracy, overall contiguity and correct scaffolding of an assembly. In order to independently asses the performance of each datatype for scaffolding the number of REAPR breaks were compared between the w2rap-only assembly (A1) and that assembly scaffolded with one datatype, namely LMP (A2), Bionano (A5) and 10x (A9).

#### Value-for-money

Cost is a huge factor in research and ultimately, impacts on decisions made regarding the technologies used. A metric was created to reflect ‘value-for-money’ by estimating the cost of each assembly and the N50 achieved. This metric is provided as N50/$1K and calculated for contig N50, scaffold N50 and the REAPR broken scaffold N50.

#### Ranking assemblies

Each assembly was ranked with regard to its performance for seven key metrics. Each assembly was given a rank-score according to its position in each metric. The top-placed assembly that performed best in a given metric, was given a rank-score of 10, the second-placed assembly was given a rank score of 9, and so on, down to the bottom-placed assembly which was given a rank-score of 1.

Assemblies were ranked for the following metrics:

1. Scaffold N50
2. REAPR broken scaffold N50
3. Contig N50
4. Percentage of genome represented by scaffolds >25 kb
5. Single-copy BUSCO orthologs
6. REAPR summary score
7. REAPR broken scaffold N50/$1K

#### Z-scores

Z-scores were used to combine scores from datasets with different means, ranges, and standard deviations and have the benefit of rewarding/penalising those assemblies with exceptionally high/low scores in any one metric. The influence of each of the seven metrics was tested by removing each metric in turn and recalculating the z-score for each assembly. These recalculations were then used to produce error bars for the final z-score figure, by providing the minimum and maximum z-score that might have occurred if any combination of six metrics was used.

## Results

### Assembly contiguity and connectivity

#### Assembly Statistics

After assembling the 10 genomes as described in Figure 1, a number of metrics were calculated for each assembly to examine contiguity and connectivity, measured by the lengths and distribution of the scaffolds within each assembly (Table 2). The mean assembly size for all genomes was 2.52Gb, slightly larger than the 2.41Gb assembly of the domestic ferret [22]. 10x-based assemblies erred on having smaller genome assembly sizes (2.46 – 2.50Gb) with the larger assemblies (2.47 – 2.66Gb) being from the PCR-free Illumina-based assemblies.

**Table 2.** Genome assembly statistics (for sequences >1kb) for all assemblies. % scores refer to percentage of scaffolds over given threshold. 47Mb is size of smallest human chromosome and hence and indication of the number of chromosome-sized scaffolds.

| No. | Assembly | No.<br>scaffolds<br>>100 kb | No.<br>scaffolds<br>>1 Mb | No.<br>scaffolds<br>>47 Mb | %<br>genome<br>>= 25kb | Longest<br>scaffold<br>(Mb) | Contig<br>N50<br>(kb) | Scaffold<br>N50<br>(Mb) | Assembly<br>Size<br>(Gb) |
| --- | --- | --- | --- | --- | --- | --- | --- | --- | --- |
| A1 | w2rap | 6,290<br>(10.7%) | 176<br>(0.3%) | 0 | 94.9 | 2.52 | 182.93 | 0.30 | 2.47 |
| A2 | w2rap +<br>lmp | 1,680<br>(4.9%) | 682<br>(2.0%) | 0 | 94.8 | 15.65 | 271.16 | 2.62 | 2.60 |
| A3 | 10x | 1,023<br>(3.9%) | 501<br>(1.9%) | 0 | 93.3 | 32.15 | 207.98 | 5.26 | 2.46 |
| A4 | 10x + lmp | 669<br>(4.2%) | 346<br>(2.2%) | 1 | 94.7 | 58.16 | 210.72 | 10.33 | 2.50 |
| A5 | w2rap +<br>bionano | 4,361<br>(7.7) | 626<br>(1.1%) | 0 | 93.8 | 6.89 | 182.93 | 0.85 | 2.66 |
| A6 | w2rap +<br>lmp +<br>bionano | 990<br>(3.0%) | 468<br>(1.4%) | 0 | 94.8 | 34.30 | 271.16 | 5.73 | 2.60 |
| A7 | 10x +<br>bionano | 604<br>(2.3%) | 336<br>(1.3%) | 0 | 97.5 | 46.79 | 207.98 | 10.84 | 2.48 |
| A8 | 10x + lmp<br>+ bionano | 409<br>(2.6%) | 218<br>(1.4%) | 7 | 97.6 | 104.38 | 210.72 | 21.01 | 2.50 |
| A9 | w2rap +<br>10x | 1,097<br>(2.4%) | 467<br>(1%) | 0 | 97.6 | 35.44 | 182.93 | 5.58 | 2.47 |
| A10 | w2rap +<br>lmp +<br>bionano +<br>10x | 447<br>(1.4%) | 235<br>(0.7%) | 3 | 97.5 | 65.13 | 271.16 | 14.05 | 2.60 |

Contig N50 for the assemblies varied between 183 kb to 271 kb. Scaffold N50 for the assemblies varied between 300 kb to 21 Mb. The increase from contig N50 to scaffold N50 varied greatly (Figure 2). The addition of LMP data to an initial short-read assembly had a varying effect. On the relatively fragmented w2rap assembly (A1), the addition of LMP reads lead to an almost 9-fold increase of the scaffold N50 but adding LMPs to the more contiguous 10x assembly (A3) resulted in a 2-fold increase. This is not unexpected as the N90 value for the 10x assembly (800kb) is 20 times greater than that of the w2rap assembly (40kb), hence the chance of mate pairs spanning the same contig and not adding to the contiguity of the assembly is much higher in the already contiguous 10x assembly. The addition of Bionano data to assemblies leads to a similar scaffold N50 increases across all assemblies, namely between a 2 and 2.8 fold increase. Finally, 10x-scaffolding data was added to scaffold assembly A1 (w2rap) and assembly A6 (w2rap + lmp + bionano). As might be expected, the effect of 10x-scaffolding data on less contiguous genomes was greater than that on more contiguous genomes. There was an 18.6-fold increase in N50 between assembly A1 (w2rap) and assembly A9 (w2rap + 10x), whereas the increase in N50 between assembly A6 (w2rap + lmp + bionano) and assembly A10 (w2rap + lmp + bionano + 10x) was less contrasting at 2.5-fold.

**Figure 2.**
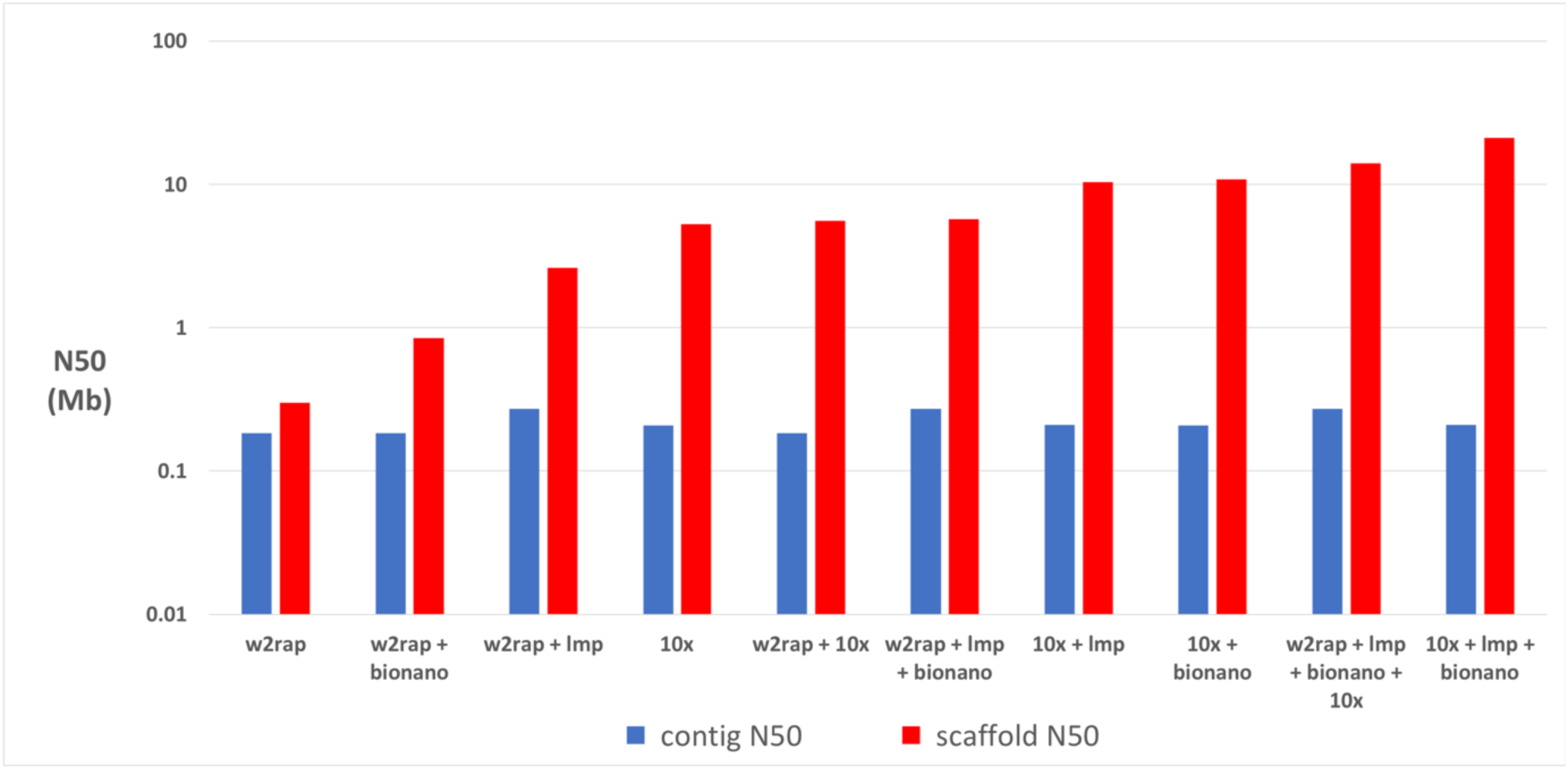
Log-scale lengths of contig N50 (blue) and scaffold N50 (red) of all ten assemblies, sorted (left to right) by scaffold N50.

Generally speaking, assemblies created with one or two data types, where one of the data types was Illumina short reads, showed the smallest increase from contig N50 to scaffold N50 (Figure 2).

#### Assembly errors and misassemblies

REAPR was used to assess the accuracy of the polecat genome assemblies by looking at low-quality regions, breakpoints (Table 3), and summary scores. (Figure 3). The percentage of error-free bases for each assembly varied between 76.05% to 85.9%. All the w2rap-based assemblies were on the low end of the scale (76.05% – 81.09%), whilst 10x-based assemblies were on the high end (84.65 % – 85.9%). Conversely, there was a trend for w2rap-based assemblies to be less affected by misassemblies (excluding those with 10x-scaffolding). Their REAPR broken N50 size reduced between 2% – 64%, whilst 10x-based assemblies reduced in N50 size between 68% – 91%. A similar pattern is seen with the number of FCD errors, where all w2rap-based assemblies (bar A10, with 10x-scaffolding) have less than 8214 FCD errors and all 10x-based assemblies have 9095 errors or more.

**Table 3.** REAPR statistics showing the percentage of error-free bases in the assembly, N50s before and after breaking at breakpoints, the percentage decrease in scaffold N50 after breaking and the fragment coverage distribution errors (FCD errors) including errors across gaps.

| No. | Assembly name | %<br>error-free | Original<br>N50 | REAPR<br>broken<br>N50 | % reduction<br>of N50 after<br>breaking | FCD<br>errors |
| --- | --- | --- | --- | --- | --- | --- |
| A1 | w2rap | 80.83 | 300334 | 294835 | 2 | 6065 |
| A2 | w2rap + Imp | 79.10 | 2616045 | 1125605 | 57 | 8213 |
| A3 | 10x | 85.90 | 5258708 | 1688764 | 68 | 11379 |
| A4 | 10x + Imp | 85.35 | 10328293 | 1860493 | 82 | 9095 |
| A5 | w2rap + bionano | 76.05 | 847448 | 521948 | 38 | 4523 |
| A6 | w2rap + Imp +<br>bionano | 78.38 | 5729516 | 2058901 | 64 | 7392 |
| A7 | 10x + bionano | 84.65 | 10843127 | 1999584 | 82 | 13068 |
| A8 | 10x + Imp + bionano | 84.75 | 21007819 | 1863460 | 91 | 11531 |
| A9 | w2rap + 10x | 81.09 | 5579606 | 573665 | 90 | 7601 |
| A10 | w2rap + Imp +<br>bionano + 10x | 77.80 | 14054335 | 1751796 | 88 | 9488 |

**Figure 3.**
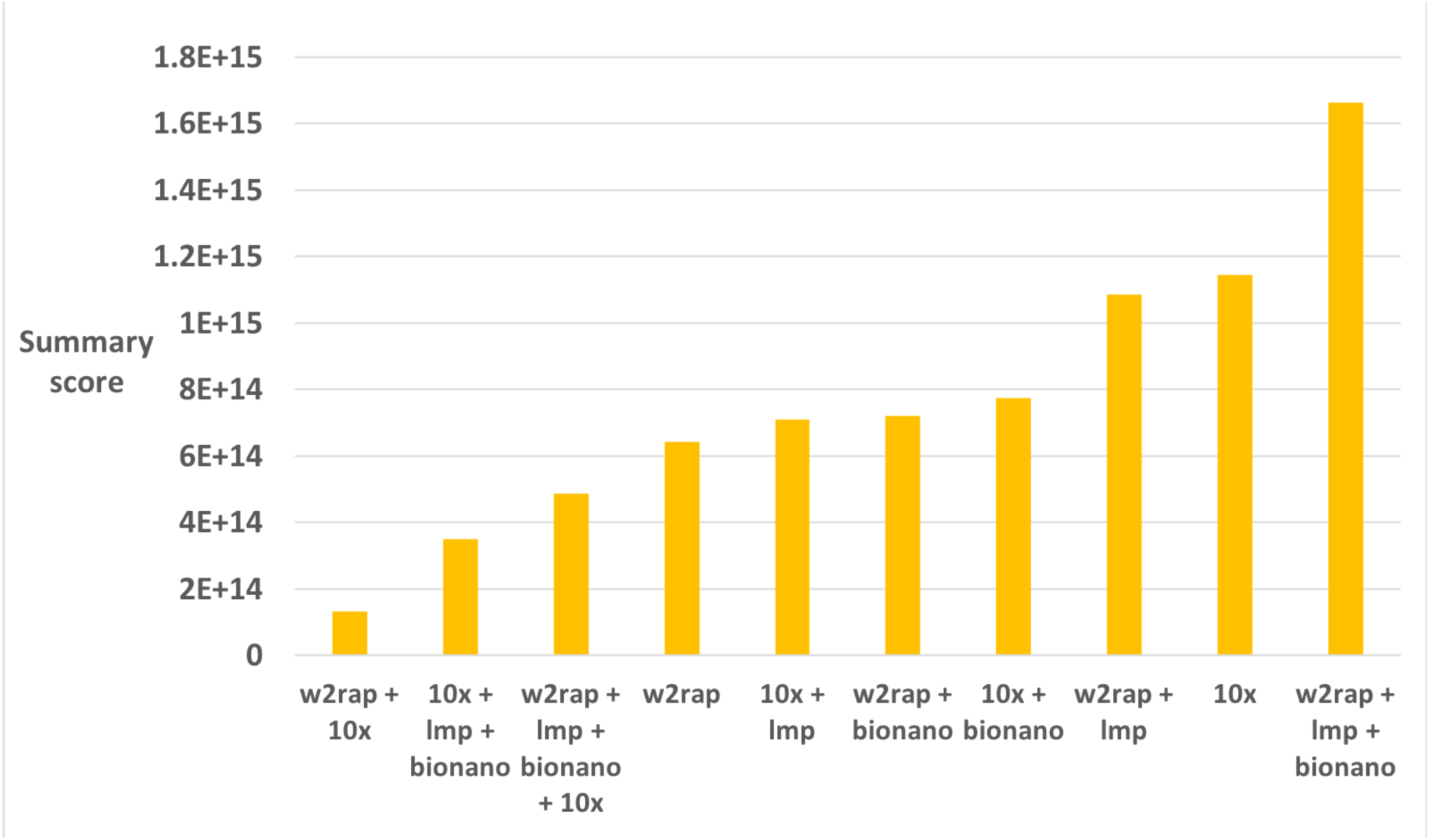
REAPR summary scores for each polecat assembly. REAPR summary scores were calculated for each assembly by multiplying the number of error-free bases with the square of the REAPR broken scaffold N50 length, and then dividing by the original scaffold N50.

Finally, the performance of each technology was independently assessed for scaffolding by comparing the number of REAPR breaks between the w2rap assembly (A1) and those scaffolded with only one datatype (LMP, Bionano, and 10x-scaffolding) (Table 4). After accounting for the 2756 breaks introduced by REAPR in the w2rap-only assembly (A1), it was found that Bionano (assembly A5) clearly performed best, containing only 729 more breaks than the original assembly (A1). Conversely, LMP (6843 more breaks) and 10x-scaffolding (7353 more breaks) datatypes had at least 9 times more breaks introduced by REAPR than Bionano. A comparison was made between the number of breaks (5252) in the 10x assembly (A3) to the 10x + LMP assembly (A4) and the 10x + Bionano assembly (A7) (Table 5). A similar pattern as above was found, with the LMP assembly having 2785 more breaks than the 10x assembly but with the Bionano assembly having only 61 more breaks, again demonstrating the accuracy of Bionano for scaffolding.

**Table 4.**
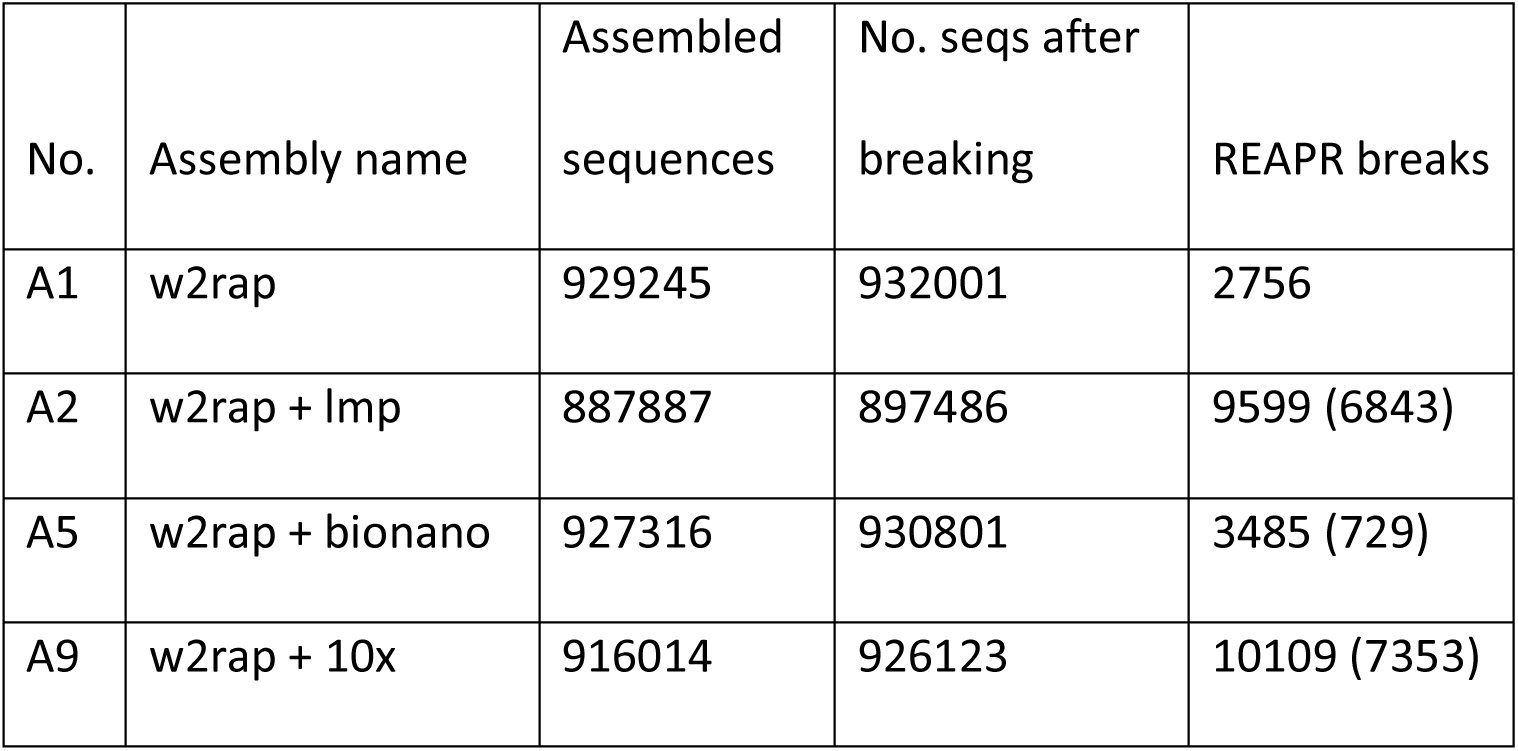
Comparison of the number of breaks introduced by REAPR for each of the technologies used to scaffold the w2rap-only assembly (A1). The number of breaks in brackets represent the number of breaks after accounting for the 2756 breaks introduce into the comparison assembly (A1).

| No. | Assembly name | Assembled sequences | No. seqs after breaking | REAPR breaks |
| --- | --- | --- | --- | --- |
| A1 | w2rap | 929245 | 932001 | 2756 |
| A2 | w2rap + Imp | 887887 | 897486 | 9599 (6843) |
| A5 | w2rap + bionano | 927316 | 930801 | 3485 (729) |
| A9 | w2rap + 10x | 916014 | 926123 | 10109 (7353) |

**Table 5.**
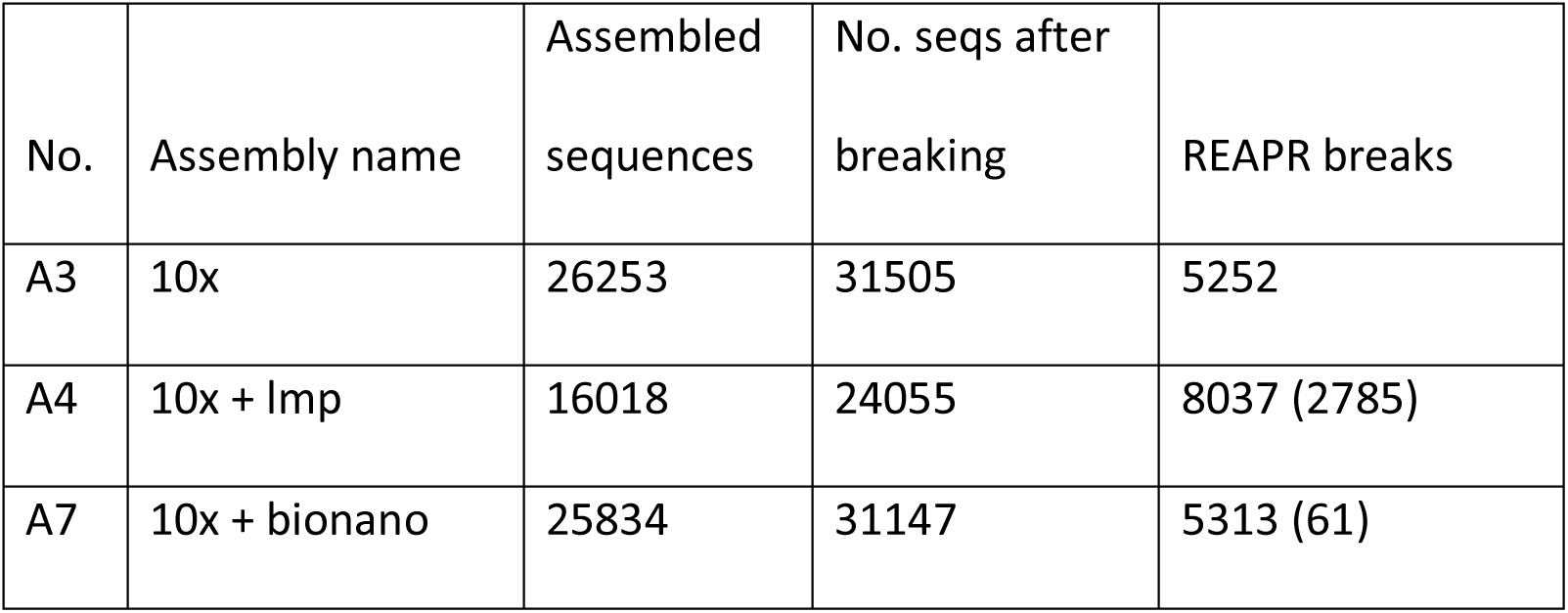
Comparison of the number of breaks introduced by REAPR for each of the technologies used to scaffold the 10x assembly (A3). The number of breaks in brackets represent the number of breaks after accounting for the 5252 breaks introduced into the comparison assembly (A3).

### Assembly Completeness

#### K-mer content

‘KAT comp’ [27] was used to compare k-mers in the Illumina PCR-free reads with k-mers in the non-Bionano assemblies (A1 – A4 and A9). ‘KAT plot’ was then used to visualise the output (Figure 4 and Supplementary Figure S1). The plots all show a similar distribution of k-mers. The black distribution at the start of the x-axis represents sequencing errors in reads and its increased width represents an increased number of errors in the reads. K-mers in these reads have not been incorporated into the final assembly. The extension of the black line along the x-axis (up to a k-mer-multiplicity of 40 on the x-axis) represents collapsed haplotypes, where k-mers from one side of a bubble in the assembly graph have been removed to construct a linear path through the graph. Any extension of the black line along the x-axis into the main red distribution (>40 k-mer-multiplicity) represents a small number of high-copy k-mers in the reads missing from the assembly. The red area in all graphs represent a normal distribution of k-mers found in the reads and occurring once in the assembly.

**Figure 4.**
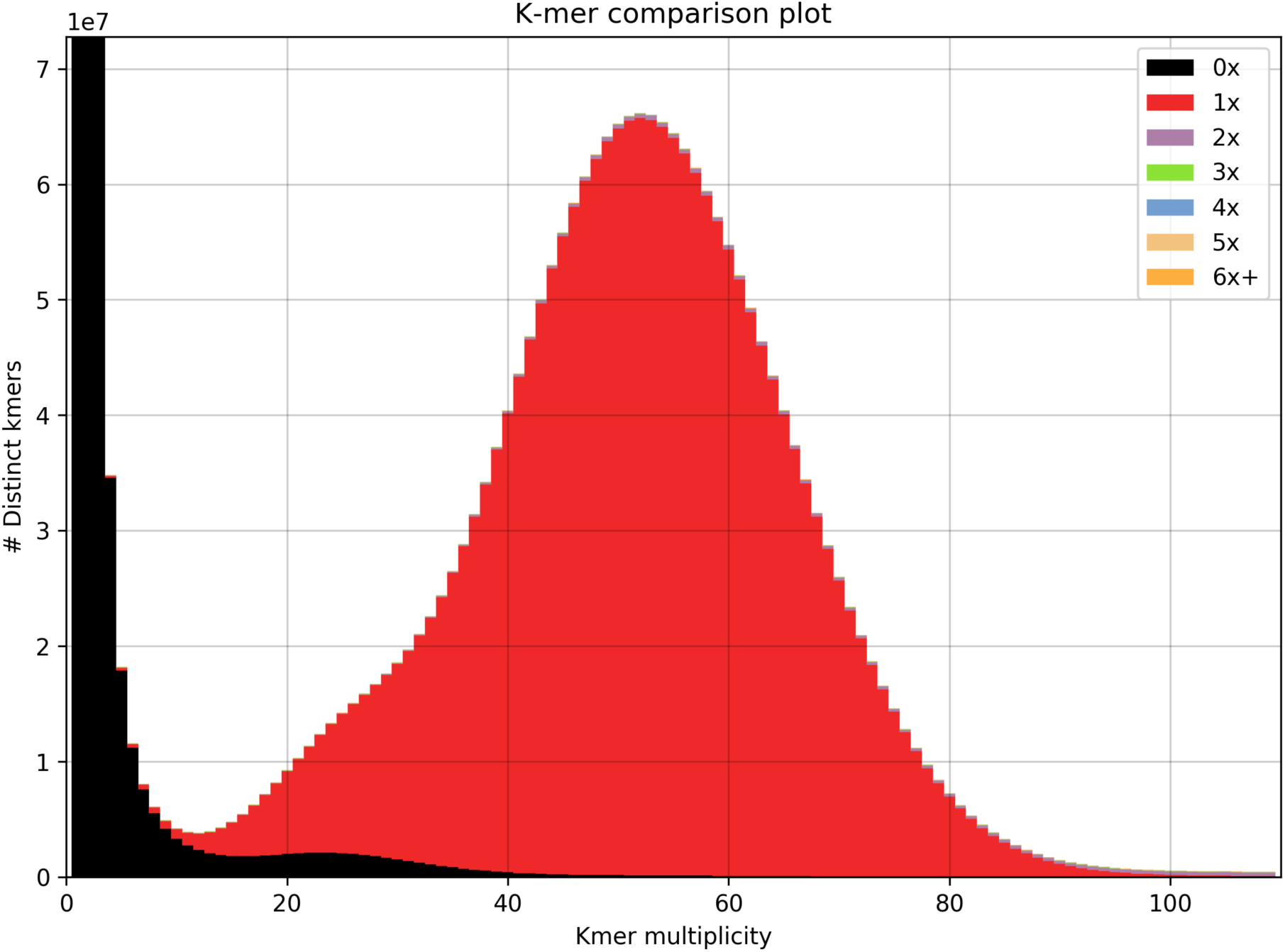
KAT k-mer plots comparing k-mer content of Illumina PCR-free reads with w2rap assembly (A1). The black area of the graphs represents the distribution of k-mers present in the reads but not in the assembly and the red area represents the distribution of k-mers present in the reads and once in the assembly.

Despite all of the assemblies being compared to the PCR-free Illumina short reads, virtually the same distribution of k-mers between the reads and assemblies was observed, showing an almost-identical distribution of k-mers from all the different read sequences and their resulting assemblies. The KAT-plots involving 10x assemblies (Supplementary Figure S1, C and D) are also characterised by some high-copy read k-mers missing from the assemblies. This suggests that the minimum size of contigs included in the final assembly (1kb) may be too high. This may also explain the slightly smaller assembly sizes obtained from the 10x-based assemblies when compared to the w2rap-based assemblies (Table 2).

#### Gene content

BUSCO was used to look at single-copy orthologs in the assemblies (Figure 5) and examine the number of single-copy, duplicated, fragmented and missing orthologs. The number of complete orthologs reconstructed varied from 3762 (92%) in Assembly A1 and 3895 (96%) in assembly A2. Of the 4077 mammalian orthologs examined 3578 (87%) were found in single copies across every assembly, 65 were missing, 31 were fragmented, and 21 duplicated across all assemblies. The w2rap assembly (A1) had the highest number of missing orthologs (117) and fragmented orthologs (198), probably down to the fragmented nature of the assembly. Adding Bionano to datasets did little to improve ortholog reconstruction (A5) and in many cases was detrimental, reducing single-copy and increasing the number of fragmented orthologs (A6, A7 and A8). Contrastingly, the addition of LMP data (A2) or 10x-scaffolding data (A9) assembly showed much improved results when compared to Bionano. Assemblies A2 (w2rap + lmp) and A9 (w2rap + 10x) both showed a significant increase in single-copy orthologs and decrease in fragmented and duplicate orthologs when compared to their single-data initial assemblies (Assemblies A1 and A3). The 10x + lmp (A4) assembly also shows improved results over the 10x-only assembly (A3), but to a lesser extent than the previous w2rap assemblies.

**Figure 5.**
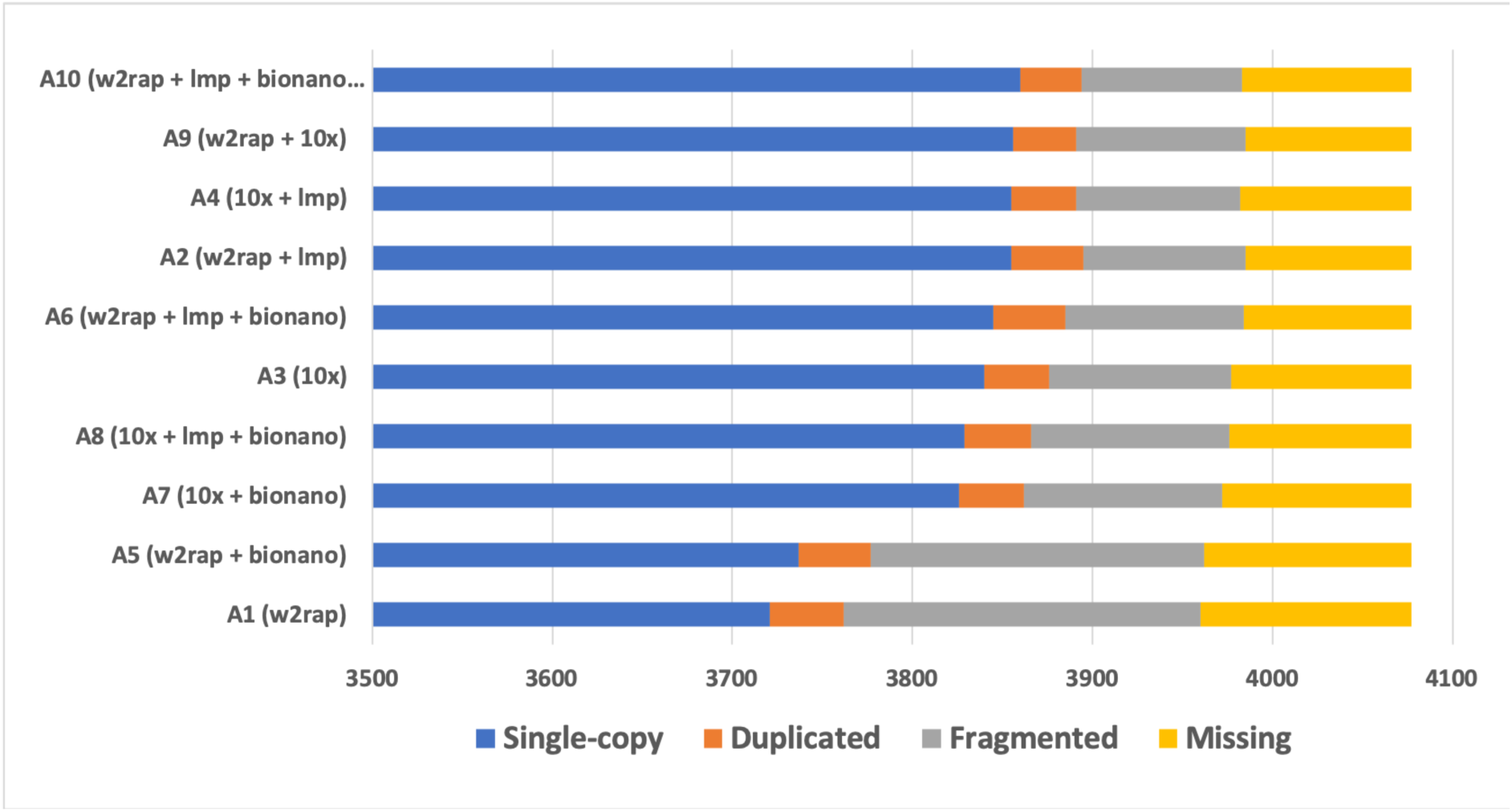
Number of single-copy (blue), duplicated (orange), fragmented (grey) and missing (yellow) orthologs from BUSCO. In order to visualise the number of duplicated, fragmented and missing orthologs, the first 3500 single-copy orthologs present in each assembly are truncated.

#### Repeats

RepeatMasker was used to look at *Carnivora*-specific repeat content in the assemblies. Long Interspersed Nuclear Elements (LINEs) and Short Interspersed Nuclear Elements (SINEs) were by far the most common classes of repeats and these are concentrated on here. A very similar picture was found between all datasets. The percentage of the genome assemblies that were masked for repeats varied between 35.82% – 39.49%, with SINEs varying between 8.4% – 9.81%. The w2rap-based assemblies were on the lower-end of both of these scales with the 10x-based assemblies on the higher-end.

A slightly different pattern was found when examining LINEs, the composition of which varied between 19.2% and 20.73%. In these repeats the w2rap-based assemblies clustered at the lower end of the scale, with the exception of assembly A1 (w2rap) and assembly A9 (w2rap + 10x), which grouped with the 10x assemblies at the higher end of the scale.

Mean divergence between each assembly and all repeat families was also calculated. It was found that the divergence between assemblies was small (24.52 – 24.60), with no defined grouping of the assemblies by datatype. This suggests an overall similar ability of each datatype to accurately reconstruct repeat sequences (Table 8)

**Table 8.** Repeat content of assemblies. % masked refers to the amount of the genome masked for all repeats, % SINEs and % LINEs reflect the percentage of the genome found to contain each of these classes, and mean divergence is calculated as ‘mismatches/(matches + mismatches)’ between queries and matches for all repeats.

| No. | Assembly name | % masked | %<br>SINEs | %<br>LINEs | mean<br>divergence |
| --- | --- | --- | --- | --- | --- |
| A1 | w2rap | 38.31 | 8.99 | 20.53 | 24.52 |
| A2 | w2rap-lmp | 37.24 | 8.75 | 19.95 | 24.56 |
| A3 | 10x | 39.49 | 9.81 | 20.73 | 24.53 |
| A4 | 10x-lmp | 39.1 | 9.71 | 20.52 | 24.54 |
| A5 | w2rap-bionano | 35.82 | 8.40 | 19.20 | 24.52 |
| A6 | w2rap-lmp-bionano | 36.79 | 8.64 | 19.71 | 24.52 |
| A7 | 10x-bionano | 39.11 | 9.72 | 20.53 | 24.60 |
| A8 | 10x-lmp-bionano | 38.89 | 9.66 | 20.41 | 24.58 |
| A9 | w2rap 10x | 38.32 | 8.99 | 20.54 | 24.55 |
| A10 | w2rap_lmp_bionano_10x | 36.79 | 8.64 | 19.70 | 24.53 |

### Value-for-money

The N50/$1K metric (see Methods) was calculated in order to provide a metric for value-for-money when considering the choice of technology and the return on money spent (Figure 6). For contig N50/$1K, the w2rap-based assemblies provide by far the best value-for-money, with the exception of those with 10x-scaffolding. Value-for-money decreases as more data is added to the w2rap assemblies. So, for contig assemblies a basic PCR-free Illumina short-read assembly provides the best value-for-money.

**Figure 6.**
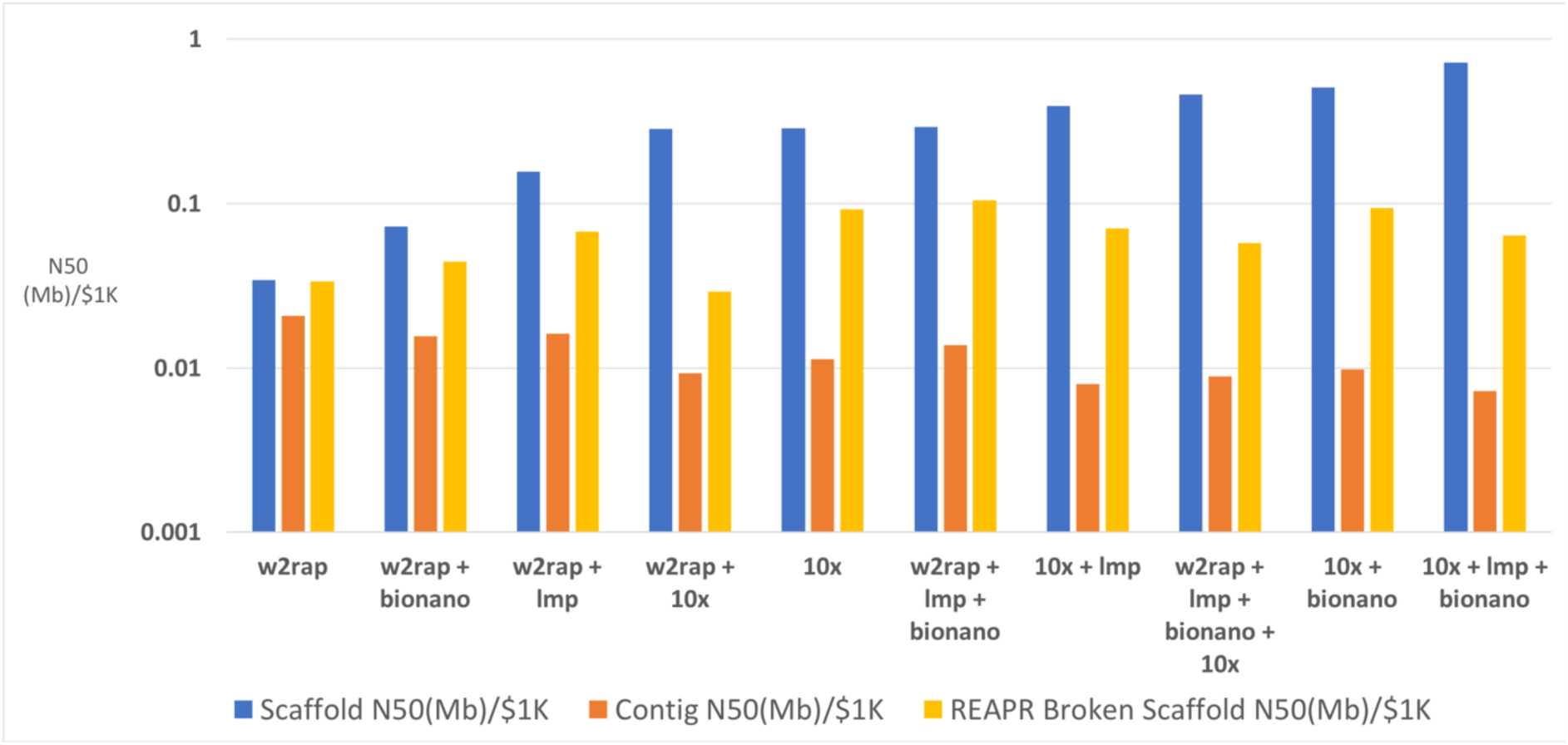
N50/$1K, providing an estimate to the cost of contig (orange), scaffold (blue) and REAPR broken scaffold (yellow) contiguity for each genome assembly. Assemblies are ranked in order of scaffold N50/$1K.

However, when looking at scaffold N50/$1K, the trend changes. Five of the six lowest-scoring assemblies constitute w2rap-based assemblies, generated with between one and three datatypes. The 10x-based assemblies show better performance when looking at scaffold N50/$1K, with three of the four highest-scoring assemblies being 10x-based. The difference in scaffold N50 between the w2rap-based and 10x-based assemblies might be expected as the short-read Illumina data does not contain the additional molecule specific linked-read information present in 10x data. Another trend is that adding more scaffolding data to a ‘base’ assembly (A1 and A3) increased the scaffold N50/$1K. Hence, adding more data to increase scaffold contiguity provides value for money, although one must judge if the amount of increase justifies the extra cost.

#### Ranking assemblies

Assemblies were ranked on a number of key metrics (see Methods), allocated a final rank-score (Supplementary Figure S2) and z-scores were calculated for each assembly (Figure 7). The order of ranking in both scoring methods agree, but the z-scores provides a better assessment of the performance of each assembly across all the metrics and not just their position in the final ranking. The general trend was that the more data included, the higher the assembly ranked. For example, the second-highest ranked assembly was A10, the only assembly with four different data types (w2rap + lmp + bionano + 10x). The highest placed assembly was assembly A6 (assembly A10, but without the final 10x scaffolding data).

**Figure 7.**
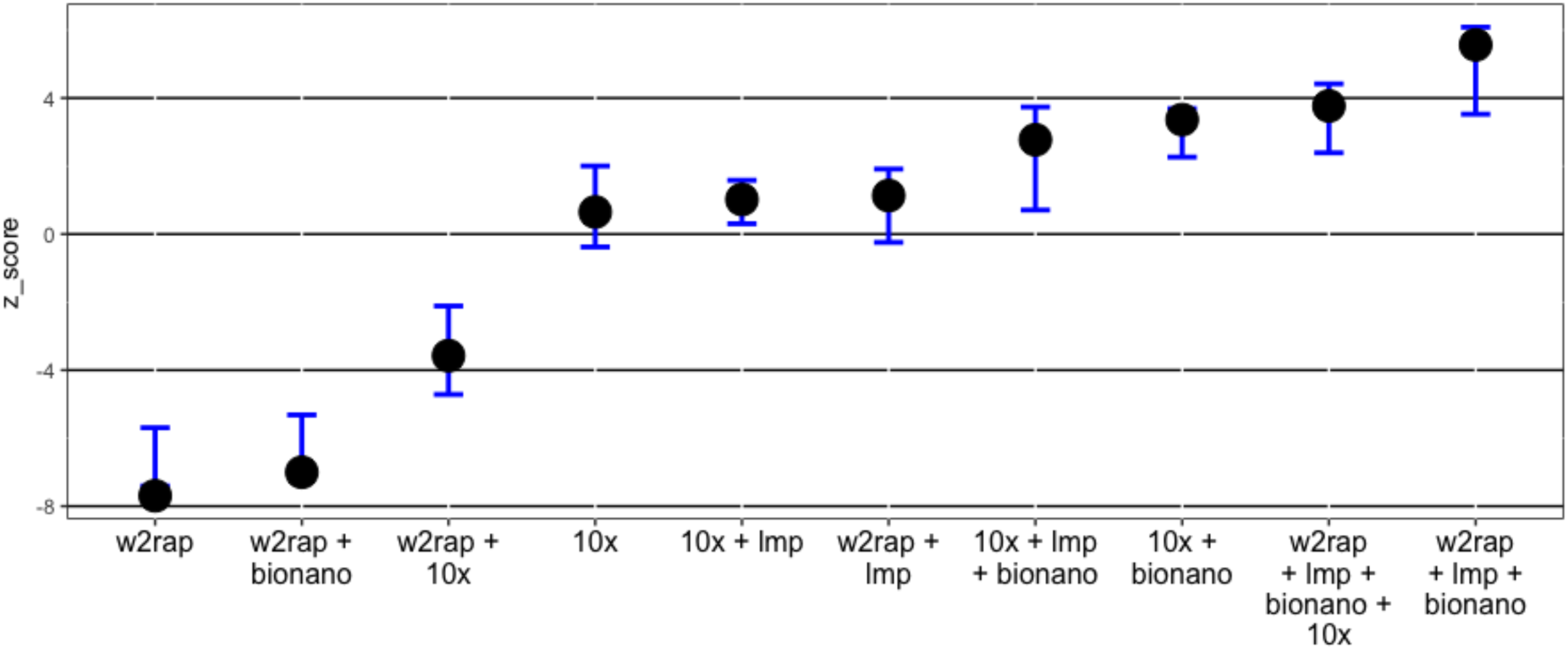
Cumulative z-scores of assemblies (solid black circles) with error bars (blue). Error bars represent the min and max cumulative z-score after removing each metric in turn and recalculating the z-score for each assembly. Wide error bars show assemblies that are strongly affected by a given metric. For example, the 10x + lmp + bionano assembly (A8) has a long lower-boundary error-bar as it has an exceptionally high scaffold N50 z-score (double that of the next nearest ranking assembly) and hence omitting this metric results in the assembly scoring much lower.

## Discussion

Although chromosome-scale assemblies are now achievable, it is often not possible or necessary to assemble the genomes of non-model organisms to such precision. A number of difficulties are faced when sequencing and assembling non-model organisms. Genome size (and as a consequence, sequencing depth), chromosome number, sequence composition and GC content are often unknown or inaccurate, the species can be highly heterozygous, and samples are often degraded. We address these factors and identify which sequencing and assembly strategies are required to answer various biological questions. PCR-free Illumina short-read, 10x Genomics linked-read, long mate paired read, and Bionano optical maps were generated from a roadkill European Polecat to create ten different genome assemblies, using different combinations of the data. The assemblies were assessed using a range of tools and ranked using seven key metrics. We find that although some genomes assemble to high contiguity, this is often at the expense of accuracy and it is often not necessary to spend additional funds on increasing contiguity to answer biological questions.

### Assembly contiguity and connectivity

As a general rule, adding more data to an assembly increases the contiguity (scaffold N50). This was observed in the assemblies here, with each assembly having a higher scaffold N50 than any ‘parent’ assembly before it. The linked reads from 10x Genomics data constantly outperform the equivalent PCR-free short-read-based assemblies, with the barcoded linked reads acting as an additional scaffolding dataset. The assemblies with the best contig N50 were those based on w2rap + LMP (namely, A2, A6 and A10). For scaffold N50 and percentage of the genome represented by scaffolds >25Kb, 10x + LMP + Bionano (A8) provided by far the best contiguity. When REAPR breaks are taken into consideration, the w2rap + LMP + Bionano assembly (A6) provides the best scaffold N50, although assembly A8, with the best initial contig N50, is still ranked third. It should be noted that Bionano and 10x (for scaffolding) added no sequence data to the assemblies and hence do not extend the contig lengths or have not connected contigs together without the need to add ‘N’s. Those assemblies scaffolded with LMPs however do increase in contig N50, reflecting previously unconnected contigs being joined without Ns.

An increase in REAPR summary scores was seen when LMP and Bionano data are added to PCR-free short read assemblies, but a decrease in summary scores when 10x-scaffolding reads are included. For 10x-based assemblies, the addition of extra data leads to a reduction in summary scores. Additionally, 10x-based assemblies tended to have more FCD errors and the breaking of assemblies at these errors affected 10x-based assemblies to a greater degree than w2rap-based assemblies. Finally, the number of breaks created by REAPR for each scaffolding technology showed that Bionano-scaffolded genomes had significantly fewer breaks than both LMPs and 10x-scaffolding. The addition of 10x-scaffolding data led to an overall reduction in summary scores suggesting that although 10x-scaffolded assemblies provide a good increase in scaffold N50, much of this increase is through misassemblies.

The increase in misassemblies with the addition of extra data is understandable. An initial, one-technology *de novo* assembly will have all the ‘easy joins’ put together and most of those will be correct. When a new datatype is added, it will have the ‘difficult joins’ to put together, making it very likely that a significant number of these will be incorrect. Bionano performs best at connecting these ‘difficult joins’

### Assembly completeness

Assembly completeness was assessed by the number of resolved BUSCO single-copy orthologs (gene content) and repeat content. Gene content was increased by adding LMP data but Bionano data tended to fragment orthologs, suggesting a degree of misassembly in the gene space. In all assemblies scaffolded with LMPs or 10x reads, the additional data was only used to scaffold the input assembly, so the introduction of extra sequence data was not responsible for the increase in ortholog reconstruction. Therefore, sequence-based scaffolding methods (LMP and 10x-scaffolding in this case) performed better in the gene space than Bionano optical mapping approach. It was considered that this might be due to the fact that domestic ferret Bionano data was used on wild polecats. However, a similar pattern was seen after scaffolding the domestic ferret genome (Ensembl MusPutFur1.0) with the Bionano data, which was obtained from the same sample used for the assembly (single copy orthologs dropped from 3857 to 3853 when scaffolded with the Bionano data).

There was a small amount of difference in repeat content between assemblies. The tendency of 10x assemblies to have a slightly higher percentage of the genome assembled as repeats probably reflects the ability of this technology to better resolve repeats than the standard short-read assemblies which collapse a large proportion of the repeats. Those repeats that were resolved in assemblies all showed a very similar divergence, regardless of the data types used.

### Value-for-money

For contig assembly, a basic PCR-free short read assembly provides the best value-for-money (A1). Adding more data does not increase the contig N50 enough to warrant the extra expense. For scaffold assembly, the story is very different. The 10x + lmp + bionano (A8) offers the best value for money. The more data added to an initial assembly, the higher the scaffold N50/$1K. When REAPR broken assemblies are taken into consideration, the 10x-based and LMP-scaffolded assemblies provide the best value, with the w2rap + lmp + bionano assembly (A6) being ranked top (Figure 6). Another feature when considering REAPR broken assemblies, is the poor performance of 10x-scaffolding (A9 and A10).

Compared to an Illumina PCR-free library, 10x Chromium libraries are expensive to produce due to higher cost of the library preparation and the additional hands-on time required associated with the protocol. This increased cost for the 10x Genomics scaffolding data and the high misassembly rate when used as a scaffolding technology, means that it scores low in this metric.

In summary, when looking at genome contiguity, if contigs are all that are required from a genome assembly, then PCR-free short-read assemblies with no additional datatypes provide the best value-for-money. If accurate scaffolds are more important, 10x data, often augmented with LMPs or Bionano provide good value-for-money, with Bionano misassembling significantly fewer scaffolds than LMPs (although LMPs perform much better in the gene space than Bionano).

### Ranking assemblies

As expected, the general trend was that the more data included, the higher the assembly ranked, although we provide evidence against the use of low-coverage 10x Genomics as a scaffolding-only technology.

### Application to non-model mammals

Although the sample quality used for non-model organisms is often sub-standard, sequencing technologies and software are still successful in assembling these samples into highly contiguous genomes. As a general rule, adding more data to an assembly increases the contiguity (scaffold N50), but the additional expense of incorporating additional data to increase contiguity is not always necessary.

For population genetics approaches, SNP-calling and large multi-species comparisons, basic short-read assemblies such as w2rap (A1) or 10x (A3) provides enough accuracy and contiguity to achieve interpretable results. 10x assemblies also have the added advantage of haplotype resolution. Where structural variation, long repeat content, gene order, or gene clusters are of importance an additional scaffolding dataset is often necessary to obtain the required precision for these analyses (A2, A4, A5, and A7), with LMP being the better data to incorporate if working in the gene space. Examples where this might be important is when dealing with gene clusters of similar genes, such as immune-related gene clusters (e.g. MHC, Interleukin, toll-like receptors, etc.). When looking at more long-range features, such as genome synteny, Bionano provides additional contiguity. Bionano though, is dependent on high-quality HMW DNA which might not be available for many organisms and appears to be the first datatype to suffer from sample degradation.

### Experimental design

Assemblies with both short contig lengths and a high number of misassemblies, can sometimes be found in very heterozygous species. Knowing the distribution of molecule lengths from a sample will provide information about the limitations of which sequencing technologies can be successfully supplied. Researchers can then design their assembly and analysis pipeline to accommodate the limitations of the sample. For example, if the molecule lengths are only in the region of 1kb, then PCR-free Illumina paired-end sequencing is the only viable option. Longer molecules, between 10 – 40 kb allow the preparation of LMP libraries and between 20 – 100 kb permit the inclusion of 10x Genomics data. Beyond that (100kb+), Bionano optical maps may also be included.

Adding long-read data, such as low-coverage PacBio or Nanopore data, will often be the only solution to overcoming complexities such as high heterozygosity or long repeats. Unfortunately, long-read data relies on high molecular weight DNA with long molecules, but as described previously, DNA samples from non-model organisms are often of low-quality and the application of these technologies may not be suitable. The quality of sample should reflect the experimental design and assembly pipeline. Development in new DNA extraction (e.g. Nanobind Magnetic Disks [32]) and sequencing technologies may provide access to low quantity and quality of DNA, which may be a potential solution to overcome the sample extraction issues.

As mentioned, with longer molecules, using long-read technologies such as PacBio and Nanopore becomes a possibility, but these require significantly more DNA (>20ng) to work successfully, as well as being associated with a much higher cost. This overcomes some of the limitations of short-read assemblies, such as characterising structural variation, sequencing through extended repetitive regions, discriminate paralogous genes and detecting disease-associated mutations, although with the drawback of requiring high-coverage due to the lower base accuracy of long read sequencing.

### Limitations of this study

In this study a combination of four different technologies have been used to create 10 different genome assemblies. An exhaustive assessment would produce many more different assemblies, so a choice of what was considered a good representation of all practical combinations was used. Additionally, different assembly software (including versions thereof) may produce slightly different results depending on the algorithms used within them. Finally, test metrics can bias results. For example, the inclusion of more cost-related metrics would bias rankings to favour cheaper assemblies, whereas more contiguity-related tests would bias results for assemblies with higher N50s. The choice of metrics was made to encapsulate genome contiguity, accuracy, error, biologically meaningful content, and cost whilst not unduly biasing the results towards any one feature of the assemblies.

### Summary

We address how different sequencing and assembly strategies are required to answer various biological questions in non-model mammals. We find that although some genomes assemble to high contiguity, this is often at the expense of accuracy and it is often not necessary to spend additional funds on increasing contiguity to answer biological questions.

Sequencing technologies and assembly software are always progressing with new sequencing chemistry releases providing longer and more accurate read sequences. Also, novel assembly algorithms promise more contiguous and accurate assemblies. Often each algorithm is dependent on the input of specific data types, with some new assembly software providing more contiguous assemblies at the expense of accuracy. It is important to fully assess the performance of an assembly by using a number of different quality assessment approaches as shown in our study, rather than relying on simple statistics such as scaffold N50, which itself can be biased by the exclusion of shorter sequences from the calculations.

Finally, given the accuracy of PCR-free assemblies and the contiguity of the 10x linked-read technology, if a PCR-free linked-read sequencing technology existed, it would provide accurate, contiguous and cheap assemblies.

## Data availability

The submission of sequencing data was brokered by the COPO platform (https://copo-project.org), funded by the BBSRC (BB/L024055/1) and supported by CyVerse UK, part of the Earlham Institute National Capability in e-Infrastructure. All datasets supporting the results of this article are available in the ENA repository under project accession number PRJEB33767.

