## Supplementary data for "Sequencing smart: *De novo* sequencing and assembly approaches for non-model mammals"

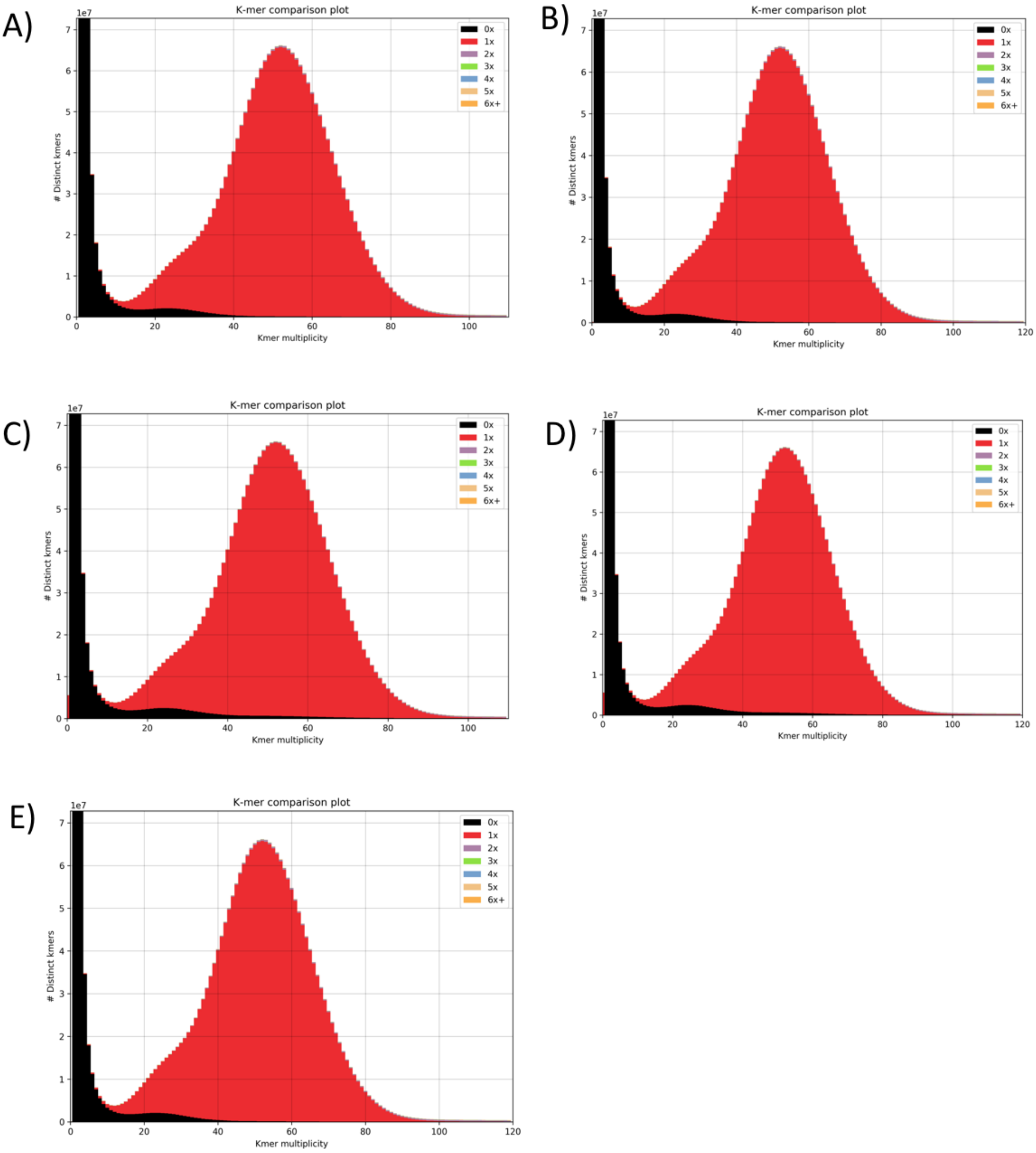


Supplementary Figure S1. KAT k-mer plots comparing k-mer content of Illumina PCR-free reads with: A) w2rap assembly (A1), B) w2rap + lmp assembly (A2), C) 10x assembly (A3), D) 10x + lmp assembly (A4), and E) w2rap + 10x assembly (A9). The black area of the graphs represents the distribution of k-mers present in the reads but not in the assembly and the read area represents the distribution of k-mers present once in the reads and once in the assembly.

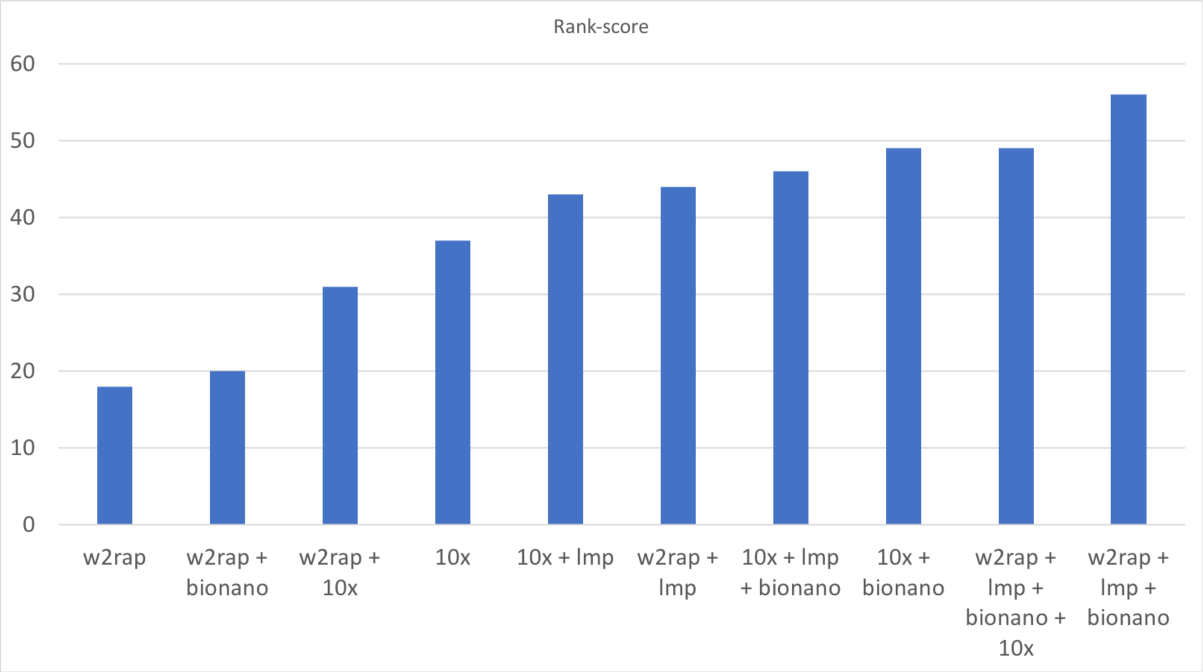


Figure S2. Each assembly was ranked over seven key metrics. A rank-score of 10 was given to the highest ranking assembly, down to a rank-score of 1 for the assembly that performed worst for the given metric.

| Assembly number | Assembly short name | Data types | Assembly strategy and software |
|---|---|---|---|
| **A1** | w2rap | PCR-free ISR | w2rap |
| **A2** | w2rap + lmp | PCR-free ISR, LMP | w2rap, SSPACE |
| **A3** | 10x | 10x Genomics | Supernova |
| **A4** | 10x + lmp | 10x Genomics, LMP | Supernova, SSPACE |
| **A5** | w2rap + bionano | PCR-free ISR, Bionano | w2rap, IrysView |
| **A6** | w2rap + lmp + bionano | PCR-free ISR, LMP, Bionano | w2rap, SSPACE, IrysView |
| **A7** | 10x + bionano | 10x Genomics, Bionano | Supernova, IrysView |
| **A8** | 10x + lmp + bionano | 10x Genomics, LMP, Bionano | Supernova, SSPACE, IrysView |
| **A9** | w2rap + 10x | PCR-free ISR, 10x Genomics | w2rap + scaff10x |
| **A10** | w2rap + lmp + bionano + 10x | PCR-free ISR, LMP, Bionano, 10x Genomics | w2rap, SSPACE, IrysView, scaff10x |

Table S1. Ten different assembly strategies using a variety of different data types: PCR-free Illumina short-read (ISR), long mate-pair (LMP), 10x Genomics Chromium library, and Bionano Genomics optical maps.
