## Supplementary methods for "Sequencing smart: *De novo* sequencing and assembly approaches for non-model mammals"

**Amplification Free Paired End Library Construction Protocol**

A total of 600ng of DNA was sheared in a 60µl volume on a Covaris S2 (Covaris, Massachusetts, USA) for 1 cycle of 40 seconds with a duty cycle of 5%, cycles per burst of 200 and intensity of 3. The fragmented molecules were then end repaired in 100µl volume using the NEB End Repair Module (NEB, Hitchin, UK) incubating the reaction at 22$^{o}$C for 30 minutes. Post incubation 58µl beads of CleanPCR beads (GC Biotech, Alphen aan den Rijn, The Netherlands) were added using a positive displacement pipette to ensure accuracy and the DNA precipitated onto the beads. They were then washed twice with 70% ethanol and the end repaired molecules eluted in 25µl Nuclease free water (Qiagen, Manchester, UK).

End repaired molecules were then A tailed in 30µl volume using in the NEB A tailing module (NEB) incubating the reaction at 37$^{o}$C for 30 minutes. To the A tailed library molecules 1µl of an appropriate Illumina TruSeq Index adapter (Illumina, San Diego, USA) was added and mixed then 31µl of Blunt/ TA ligase (NEB) added and incubated at 22$^{o}$C for 10m. Post incubation 5µl of stop ligation was added and then the reaction incubated at room temperature for 5 minutes. Following this incubation 67µl beads of CleanPCR beads (GC Biotech, Alphen aan den Rijn, The Netherlands) were added and the DNA precipitated onto the beads. They were then washed twice with 70% ethanol and the end repaired molecules eluted in 100µl nuclease free water. Two further CleanPCR bead based purifications were undertaken to remove any adapter dimer molecules that may have formed during the adapter ligation step. The first with 0.9x volume beads, the second with 0.6x and the final library eluted in 25µl Resuspension Buffer (Illumina).

Library QC was performed by running a 1µl aliquot on a High Sensitivity BioAnalyser chip (Agilent, Stockport, UK) and the DNA concentration measured using the High Sensitivity Qubit (Thermo Fisher, Cambridge, UK). To determine the number of viable library molecules the library was subjected to quantification by the Kappa qPCR Illumina quantification kit (Kapa Biosystems, London, UK) and a test lane run at 10pM on a MiSeq (Illumina) with 2x300bp reads to allow the library to be characterised prior to generation of the 60x coverage required on the Hiseq2500s (Illumina) with a 2x250bp read metric.

**Long Mate Pair Library Construction Protocol**

For the Tagmentation reactions 3µg and 6ug of Genomic DNA was prepared in 308µl and then 80µl 5x Tagment Buffer Mate Pair (Illumina) added followed by 12µl Mate Pair Tagmentation Enzyme (Illumina) and the reaction gently vortexed to mix. This was then incubated for 30 minutes at 55$^{o}$C, 100µl of Neutralize Tagment Buffer (Illumina) added and then incubated at room temperature for 5 minutes. A 1x volume bead clean-up was performed with CleanPCR beads and the DNA eluted in 165µl of Nuclease free Water . A 1µl aliquot was run on a BioAnalyser 1200 chip and DNA concentration determined using a Qubit HS Assay.

Strand Displacement was performed by combining 162µl of tagmented DNA, 20µl 10x Strand Displacement Buffer (Illumina), 8µl dNTPs (Illumina) and 10µl Strand Displacement Polymerase (Illumina). This was then incubated at room temperature for 30 minutes. A 0.75x volume bead clean-up was performed with CleanPCR beads and the DNA eluted in 16µl of Nuclease free Water and the eluted DNA from the 3µg and 6ug reactions pooled. A 1µl aliquot was diluted 1:6 and run on a BioAnalyser 1200 chip and DNA concentration determined using a Qubit HS Assay.

Size selection was performed on a Sage Science ELF (Sage Science, Beverly, USA). The 30µl in each of collection wells was replaced with fresh buffer and the collection and elution current checked prior to loading the sample. To 30µl of the pooled Strand Displaced reaction 10µl of loading solution was added and then loaded onto a 0.75% Cassette which was configured to separate the sample for 3 hours 30 minutes and then eluting each fraction for 35 minutes. Post size selection, the 30µl from each of the 12 collection wells was recovered and the DNA concentration determined using a Qubit HS Assay.

Circularisation was performed by combining 30µl of size fractionated DNA, 12.5µl of 10x circularisation buffer (Illumina), 3µl Circularisation Enzyme (Illumina) and 85µl nuclease free water. These were then incubated at 30$^{o}$C overnight. Linear DNA was digested by adding 3.75µl Exonuclease (Illumina) and incubating at 37$^{o}$C for 30 minutes followed by 70$^{o}$C for 30 minutes to denature the enzyme and 5µl of stop ligation (Illumina) added. During exonuclease treatment 240µl of M280 Dynabeads (Thermo Fisher) were prepared by washing twice with 600µl Bead Bind Buffer (Illumina) before resuspending in 1560µl Bead Bind Buffer. Circularised DNA was then sheared in a 130µl volume on a Covaris S2 for 2 cycles of 37seconds with a duty cycle of 10%, cycles per burst of 200 and intensity of 4.

To 130µl fragmented DNA 130µl of washed M280 beads was added, mixed and then placed on a lab rotator at room temperature for 20 minutes. Library molecules bound to M280 beads were then washed four times with 200µl Bead Washer Buffer (Illumina) and twice with 200µl Resuspension Buffer (Illumina).

A master mix containing 1105µl nuclease free water, 130µl 10x End Repair Reaction Buffer (NEB, Hitchin, UK) and 65µl end repair enzyme mix (NEB) was prepared and 100µl added to each tube, mixed with the beads and incubated at room temperature for 30 minutes. End repaired library molecules bound to M280 beads were then washed four times with 200µl Bead Washer Buffer and twice with 200µl Resuspension Buffer.

A master mix containing 325µl nuclease free water, 39µl A Tailing 10x Reaction Buffer (NEB) and 26µl A tailing enzyme mix (NEB) was prepared and 30µl added to each tube, mixed with the beads and incubated at 37$^{O}$C for 30 minutes. To the A tailed library molecules 1µl of the appropriate Illumina Index adapter (Illumina) was added and mixed then 31µl of Blunt/ TA ligase (NEB) added and incubated at room temperature for 10m. Post incubation 5µl of stop ligation added and then the adapter ligated library molecules bound to M280 beads were then washed four times with 200µl Bead Washer Buffer and twice with 200µl Resuspension Buffer.

A master mix containing 240µl nuclease free water, 300µl 2x Kappa HiFi (Kappa Biosystems) and 60µl Illumina Primer Cocktail (Illumina) was prepared and 50µl added to each tube, mixed with the beads and the contents, including beads, transferred to a 200µl PCR tube. Each sample was then subjected to amplification on a Veriti Thermal Cycler (Thermo Fisher) with the following conditions:- 98°C for 3 minutes, 8, 10 or 12 cycles of PCR depending upon copy number entering circularisation of 98°C for 10 seconds, 60°C for 30 seconds, 72°C for 30 seconds followed by 72°C for 5 minutes and Hold at 4°C.

Post amplification the PCR tubes were placed on a magnetic plate, the beads allowed to pellet and then 45µl of the PCR transferred to a 2ml Lobind Eppendorf Tube. To this 31.5µl beads of CleanPCR beads were added to precipitate the DNA, the beads washed twice with 70% ethanol and the final library eluted in 20µl resuspension buffer. Library QC was performed by running a 1µl aliquot on a High Sensitivity BioAnalyser chip and the DNA concentration measured using the High Sensitivity Qubit. Libraries to be sequence were then pooled based on DNA concentration and the quantification of the pool was determined by the Kappa qPCR Illumina quantification kit with the pool run at 10pM on a HiSeq with a 2x250bp reads read metric.

Reads generated were then processed through NextClip which takes LMP FASTA reads and looks to categorise them into four groups based on the presence of the Nextera adapter junction sequence. Category A pairs contain the adaptor in both reads, Category B pairs contain the adaptor in only read 2, Category C pairs contain the adaptor in only read 1, Category D pairs do not contain the adaptor in either read. NextClip also uses a k-mer-based approach to estimate the PCR duplication rate while reads are examined.

**10x Genomics Library Construction**

Polecat DNA was diluted down from 4.18ng/ul to 1ng/ul using Qiagen EB and checked by QuBit 2.0 Fluorometer (Thermo-Fisher). This was diluted in half as it was denatured. Finally, 2.5ul of the diluted and denatured DNA was loaded onto the 10X chip which equated to 1.25ng.

Gel beads, oil and master mix were loaded into the appropriate wells on the 10X chip. The DNA was loaded along with the master mix. The chip was then loaded into the 10X Genomics Chromium instrument.

The formed emulsion was removed and placed into the thermocycler for 3 hours at 30°C. GEMs were cleaned up using recovery reagent (10X Genomics), Silane beads (Thermo Fisher) and Tween-20 (Sigma-Aldrich) as per manufacturer's instructions. DNA is cleaned up using 0.7X ratio of SPRISelect beads (Beckman Coulter).

The library underwent simultaneous End Repair and A tailing on the thermocycler at 20°C for 30 minutes and then 65°C for 30 minutes. Adapters were then ligated at 20°C for 15 minutes. A 0.8X ratio SPRISelect clean-up was performed. Sample Indexes were added during the final 8 cycle PCR amplification.

| Temperature | Time |
|---|---|
| **98°C (step 1)** | 45 secs |
| **98°C (step 2)** | 20 secs |
| **54°C (step 3)** | 30 secs |
| **72°C (step 4)** | 20 secs |
| **Steps 2-4**<br>**for 7 or 8 cycles in total** | |

| | |
|---|---|
| **72°C (step 5)** | 60 secs |
| **4°C (step 6)** | Hold |

A dual-SPRI size selection was performed at 0.5X then 0.7X ratios.

Library sizes were checked using the High Sense DNA bioanalyzer (Agilent). Molarity was checked using qPCR (KAPA Library Quant kit (Illumina), ABI Prism qPCR Mix, Kapa Biosystems).

The library was clustered on the flow cell using the qPCR molarity. It was clustered at 8pM along with 1% PhiX spike-in (Illumina). Two lanes of a Rapid Run v2 flow cell were run on an Illumina HiSeq2500. 150bp PE, 8bp i7 index read, 0bp i5 index read. The libraries clustered at around 1100K/$mm^2$.

Manufacturer's instructions:

https://assets.contentful.com/an68im79xiti/4z5JA3C67KOyCE2ucacCM6/d05ce5fa3dc4282f3da5ae7296f2645b/CG00022_GenomeReagentKitUserGuide_RevC.pdf

**Bionano**

The Bionano genome maps were prepared by taking approxmately 10mg 10mg of Polecat from the sample stored in 100% EtOH. The IrysPrep Animal Tissue DNA Isolation from Fibrous Tissue protocol was followed using the IrysPrep Animal Tissue DNA kit (RE-013-10) from Bionano Genomics. The Polecat sample was cut up into <3mm pieces for homogenisation and then fixed in a 2% formaldehyde solution in kit-provided Homogenisation Buffer (HB) for 30 minutes on ice before blending using a Qiagen Tissueruptor. Spin at 1500x g for 5 mins in centrifuge, remove supernatant and re-suspend in around 50 µl of HB buffer using wide bore tip to a total volume of 66µl. 40µl of LMP agarose was then melted at 70°C and cooled to 43°C before addition with the cell resuspension and mixed using a wide bore tip. One plug of around 90ul was cast using the Chef Mammalian Genomic DNA Plug Kit (Bio-Rad 170-3591). Once set at 4°C the plug was added to a lysis solution containing 200 µl proteinase K (QIAGEN 158920) and 2.5ml of Bionano lysis Buffer. This was put at 50°C for 2 hours on thermomixer, making a fresh proteinase K solution to incubate overnight. The 50ml tubes were then removed from the thermomixer for 5 minutes before 50µl RNAse A (Qiagen158924) was added and to the tubes, returned to the thermomixer for a further hour at 37°C. The plugs were then washed 7 times in Wash Buffer supplied in Chef kit and 7 times in 1xTE. The plug was removed and melted for 2 minutes at 70°C followed by 5 minutes at 43°C before adding 10ul of 0.2U/µl of GELase (Cambio Ltd G31200). After 45 minutes at 43°C the melted plug was dialysed on a 0.1uM membrane (Millipore VCWP04700) sitting on 15ml of 1xTE in a small petri dish. After 45 minutes the sample was removed with a wide bore tip and mixed gently 5 times and left overnight at 4°C. A small amount was removed to QC on an Opgen Argus Q-Card and Qubit HS for the DNA concentration. 300ng of DNA was taken into the NLRS (Nick, Label, Repair and Stain) reaction using 1 µl Nt.BspQI (NEB R0644S). The optical maps were then generated using the Bionano Irys platform.
